# SPOP Promotes Ubiquitination and Degradation of MyD88 to Suppress the Innate Immune Response

**DOI:** 10.1101/831586

**Authors:** Qinghe Li, Fei Wang, Qiao Wang, Maiqing Zheng, Ranran Liu, Huanxian Cui, Jie Wen, Guiping Zhao

**Affiliations:** Institute of Animal Sciences, Chinese Academy of Agricultural Sciences, Beijing, China

**Author notes:** The authors declare no competing financial interests. These authors contributed equally: Qinghe Li, Fei Wang. Correspondence and requests for materials should be addressed to G.Z.

## Abstract

As a canonical adaptor for Toll-like receptor (TLR) family, MyD88 has crucial roles in host defence against infection of microbial pathogens and its dysregulation might induce autoimmune diseases. Here we demonstrate that the Cullin 3-based ubiquitin ligase adaptor SPOP recognizes the intermediate domain and degrades chMyD88 through the proteasome pathway. Knockdown or genetic ablation of chSPOP leads to aberrant elevation of the chMyD88 protein. Consequently, ChSPOP negatively regulates the activity of NF-κB pathway and thus the production of IL-1β and IL-8 upon LPS challenge. Furthermore, SPOP deficiency mice are more susceptible to infection of *Salmonella typhimurium*. Collectively, these findings demonstrate chMyD88 as a bona fide substrate of chSPOP and uncover a mechanism by which chSPOP suppresses the innate immune signaling.

**Author Summary:** MyD88 is a central adaptor mediating the initiate of innate immune response and production of proinflammatory cytokines that restrain pathogens and activate adaptive immunity. Although MyD88 is crucial for the host to prevent pathogenic infection, misregulation of MyD88 abundance might lead to autoimmune diseases. Thus, degradation of MyD88 is a canonical mechanism to terminate cytokines production. Here we characterized a novel E3 ligase SPOP that target MyD88 for degradation. SPOP attenuated IL1β and IL8 production through K48-linked polyubiquitination and degradation of MyD88 and thus impaired immune responses. SPOP deficient mice show more susceptibility to infection by *Salmonella typhimurium*. These findings demonstrate that SPOP is a negative regulator of MyD88-dependent pathway activation triggered by LPS and *Salmonella typhimurium*, which helps the host to maintain immune homeostasis.

## Introduction

The host innate immune system is the first line of defense against invading pathogens and relies on efficient recognition of microbial agents. Activation of the host innate immune response requires the detection of pathogen-associated molecular patterns (PAMPs), including proteins, lipids, carbohydrates and nucleic acids[1]. Pathogen-associated molecular patterns are recognized by pattern recognition receptors such as Toll-like receptors (TLRs), NOD-like receptors, retinoic acid-inducible gene 1 (RIG I)-like receptors and C-type lectin receptors[2, 3]. Toll-like receptors are the most widely used pattern-recognition receptors that are involved in the recognition of PAMPs including bacterial lipopolysaccharides (LPSs), flagellins, fungi and viral nucleic acids[4]. Upon recognition of the PAMPs, TLRs recruit downstream adaptors such as MyD88 to activate intracellular signaling pathways that result in the production of interferons and proinflammatory cytokines in mammals to antagonize the infection of pathogens [4, 5].

Toll-like receptors are widespread and have been found in both animal and plant phyla, indicating that these receptors were part of an ancient pathogen surveillance system[6]. Most orthologues of the TLRs in human have been identified and show similar functions with their mammalian counterparts in chickens, such as TLR2–TLR8. Like mammals, pattern recognition of pathogens by TLRs has crucial effects on the activation of innate immune responses in chickens. In chickens, TLR2 recognizes peptidoglycan, TLR4 binds LPS and TLR5 senses flagellin, and these mechanisms are almost the same as in mammals[7]. However, the TLR repertoire is unique in chickens. TLR9, which is responsible for sensing CpG DNA, is absent in chickens. Instead chickens utilize TLR21 to recognize the unmethylated CpG DNA motifs commonly found in bacteria [8].

As a central adaptor for TLR signaling, MyD88 converts signals from TLRs to activate downstream pathways. MyD88 is subjected to many protein modifications, such as phosphorylation and ubiquitination[9–12]. Mutation of the PTPN6 gene that encodes the protein tyrosine phosphatase Src homology region 2 domain-containing phosphatase-1 has been linked with autoinflammatory and autoimmune diseases, and phosphorylation of MyD88 at tyrosine residues 180 and 278 by spleen tyrosine kinase, which is suppressed by SHP1, is a prerequisite for the induction of inflammatory disease in PTPN6-mutated mice [9]. Protein polyubiquitination and deubiquitination have been shown to play critical regulatory roles in host innate immunity. Previous studies identified E3 ligases that modulated TLR signaling by promoting the polyubiquitination and degradation of MyD88, including Nrdp1, Smurf and Cbl-b[10–12]. Phosphorylation of OTUD4 confers K63 deubiquitinase activity to deubiquitinate MyD88 and subsequently deactivate TLR-mediated NF-κB signaling [13].

Ubiquitin is a small polypeptide that is covalently added into proteins by ubiquitin ligase complexes, and the targeted substrate then undergoes proteasome-dependent protein degradation[14]. The substrate specificity for ubiquitin ligation is dependent on the E3 ligase, which recruits substrate by direct protein interaction. SPOP is a protein that acts as an adaptor for the Cul3-RBX1 E3 ubiquitin ligase complex, and SPOP selectively binds to its substrates via the N-terminal domain [15]. SPOP has been shown to be linked to the ubiquitination and degradation of a number of proteins in Drosophila and humans, including AR, DAXX, SENP7, Ci/Gli and macroH2A[16–20]. Genome-wide mutation analyses have revealed the high mutation frequency of SPOP, with mutations predominantly occurring in its substrate recognition MATH domain in many cancer types, such as prostate and kidney cancer [21]. Previous studies have shown that SPOP plays important roles in tumorigenesis, cell apoptosis, X chromosome inactivation and animal development[17–19, 22]; however, the association between SPOP and host innate immunity remains poorly understood.

In this study, we identified SPOP as the ubiquitin ligase adaptor that directly promotes K48-linked polyubiquitylation and destabilizes the MyD88 protein. We also demonstrated that the SPOP is critical for regulating NF-κB signaling and innate immune response to *Salmonella* infection.

## Results

### SPOP interacts and colocalizes with MyD88

Since protein ubiquitination has emerged as an important regulatory mechanism for MyD88 signaling, we investigated whether there are other E3 ubiquitin ligases involved in the regulation of MyD88. In the amino acid sequence of chMyD88, we noticed canonical S/T-rich motifs that are the binding consensus amino acid motif of the SPOP-Cul3-Rbx1 E3 ligase complex[16]. We therefore constructed expression vectors of chMyD88 and chSPOP and transfected them into chicken embryonic fibroblasts (DF1 cells) to investigate the association between chMyD88 and chSPOP. As expected, exogenously introduced chMyD88 interacted with chSPOP *in vivo* (Fig 1A). The same interaction was observed between human and mouse MyD88 and SPOP in human cervical carcinoma cells (Hela cells) and chinese hamster ovary cells (CHO cells) (S1A and S1B Fig). We also utilized an antibody against chSPOP and our results demonstrated that endogenous chMyD88 could co-immunoprecipitate with chSPOP (Fig 1B). Consistently, immunofluorescence analysis also demonstrated the colocalization between MyD88 and SPOP (Fig 1C). Taken together, these data suggest that SPOP could interact and co-localize with MyD88.

**Fig 1.**
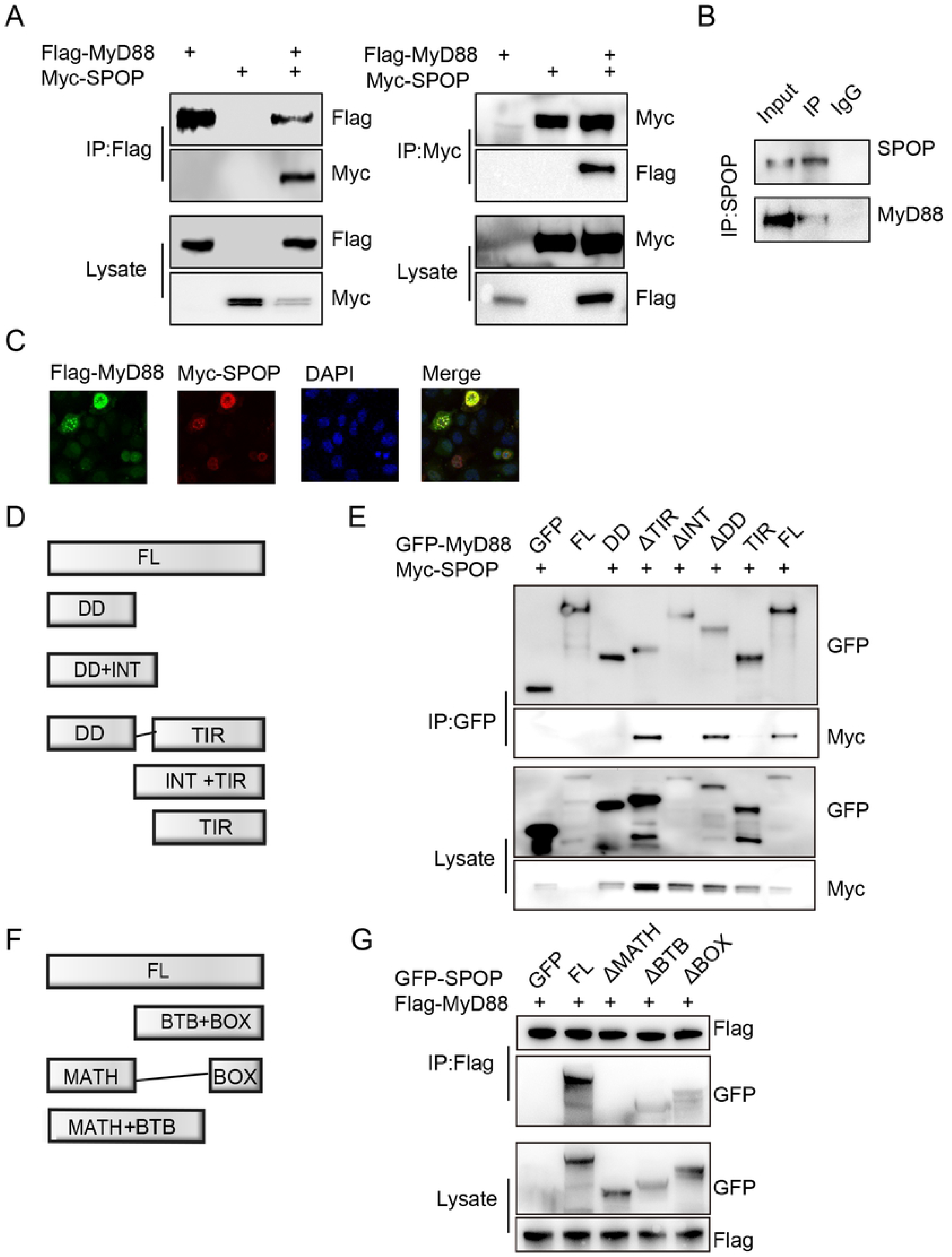
Interaction of chMyD88 with chSPOP. (A) Chicken DF1 cells were transfected with indicated plasmids. Immunoprecipitation were carried out to detect the interaction between chMyD88 and chSPOP by using the anti-FLAG or anti-MYC antibody, followed by immunoblot analysis with indicated antibodies. (B) Co-immunoprecipitation of endogenous chSPOP with endogenous chMyD88. Cell lysates were immunoprecipitated by anti-SPOP or control IgG antibody, followed by immunoblot with indicated antibodies. (C) Chicken DF1 cells transfected with Flag-tagged chMyD88 and Myc-tagged chSPOP. Then cells were fixed and incubated with anti-Flag and anti-Myc, followed by incubation with secondary antibody. Nuclei were stained with DAPI. The colocalization between chMyD88 and chSPOP was detected by confocal microscopy (D) Schematic presentation of chMyD88 and its truncation mutants. FL, full length. DD, death domain. INT, intermediate domain. TIR, Toll Toll/interleukin-1 receptor homology (TIR) domain. (E) GFP-tagged chMyD88 or its truncated mutants and Myc-tagged chSPOP were individually transfected into chicken DF1 cells. Cell lysates were immunoprecipitated with anti-GFP antibody and then immunoblotted with indicted antibodies. (F) Schematic diagram of chSPOP and its truncation mutants. MATH, meprin and TRAF homology (MATH) domain. BTB, bric-a-brac, tramtrack and broad complex/POZ domain. BOX, 3-box domain together with the C-terminal nuclear localization sequence. (G) GFP-tagged chSPOP or its truncated mutants and FLAG-tagged chMyD88 were individually transfected into chicken DF1 cells. Cell lysates were immunoprecipitated with anti-FLAG antibody and then immunoblotted with indicted antibodies.

MyD88 contains an N-terminal death-like domain, an intermediate domain and a C-terminal Toll/interleukin-1 receptor homology domain[23]. To further map the protein domain of chMyD88 that mediates the interaction with chSPOP, we constructed a series of GFP-tagged full-length and truncated chMyD88 mutants and analyzed the interactions between the full-length and truncated chMyD88 constructs with Myc-tagged recombinant full-length chSPOP (Fig 1D). We found that chSPOP co-precipitated with wild-type chMyD88, the death-like domain truncated chMyD88 mutant and the TIR domain truncated chMyD88 mutant, but not with the intermediate domain truncated chMyD88 mutant (Fig 1E), indicating that chMyD88 interacted with chSPOP via its intermediate domain. Previous studies have reported that SPOP comprised an N-terminal meprin and TRAF homology domain, a bric-a-brac, tramtrack and broad complex (BTB)/POZ domain, and a 3-box domain together with the C-terminal nuclear localization sequence[24]. Full-length or truncated forms of chSPOP were co-transfected with chMyD88 into chicken cells and co-immunoprecipitation assays showed that only the MATH domain of chSPOP was required for the interaction with chMyD88(Fig 1F and 1G), which was consistent with the finding that the MATH domain of chSPOP was primarily involved in substrate recognition and binding[15].

### chSPOP promotes proteasomal degradation of chMyD88

We next examined whether chMyD88 was subject to chSPOP-mediated protein degradation. As expected, exogenously expressed chSPOP efficiently decreased the expression of chMyD88 in a dose-dependent manner (Fig 2A). However, the mRNA level of chMyD88 remained unchanged when the expression of chSPOP was altered (S2A and S2B Fig), indicating that chSPOP regulated chMyD88 at the translational rather than the transcriptional level. Consistent with this finding, knockdown of endogenous chSPOP led to an increase in chMyD88 abundance (Fig 2B and Fig S3). The observed decease in chMyD88 by chSPOP was rescued by the proteasome inhibitor MG132 (Fig 2C), indicating that chSPOP promoted the degradation of chMyD88 in an ubiquitin-proteasome-dependent way.

**Fig 2.**
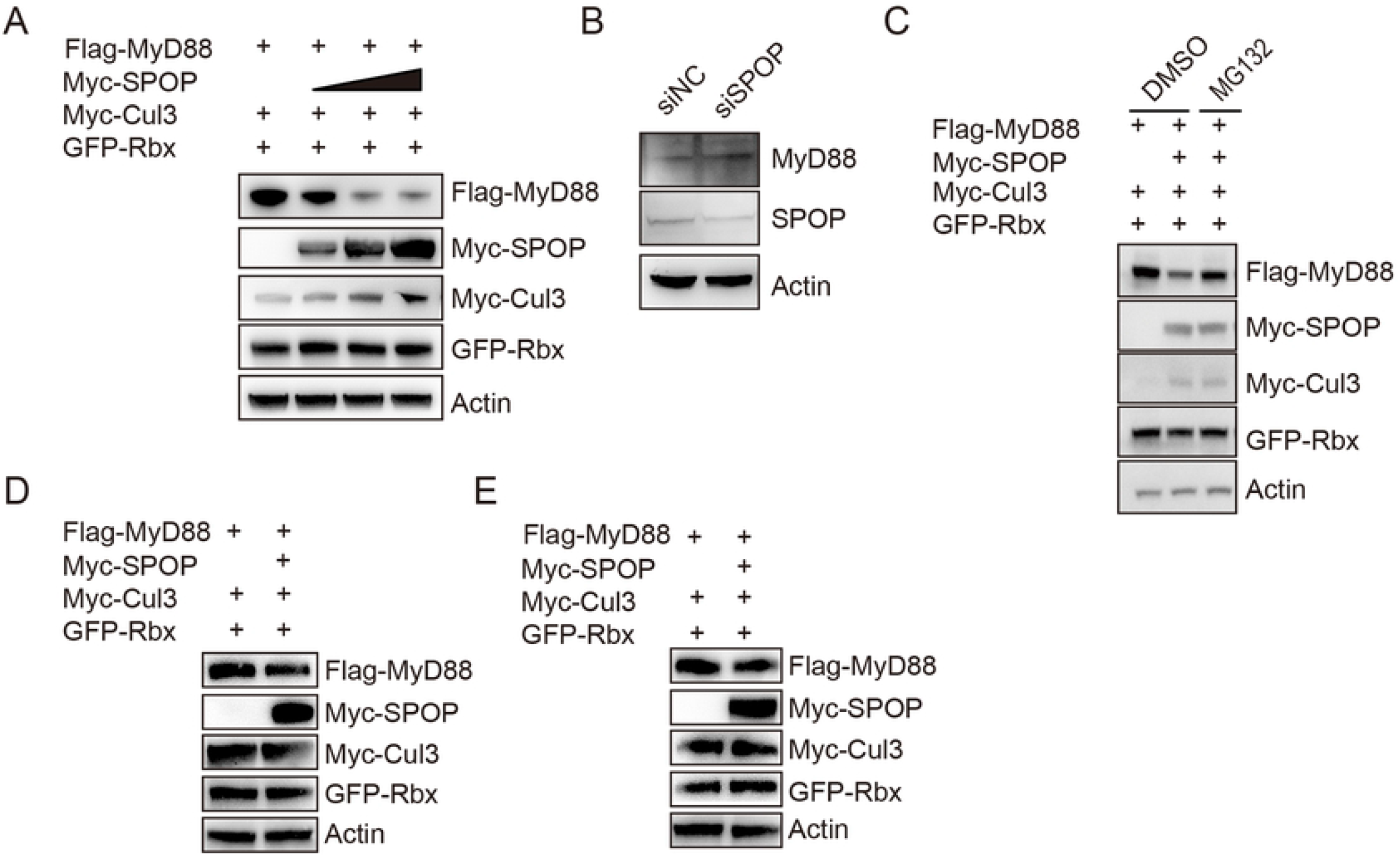
ChSPOP promotes proteasomal degradation of chMyD88. (A) Immunoblot analysis of chMyD88 in cell lysates of chicken DF1 cells transfected with chCUL3, chRBX1 and increasing doses of Myc-tagged chSPOP (0, 0.4, 0.8, 1.6μg). (B) Immunoblot analysis of endogenous chMyD88 in chSPOP inhibited cells by siRNA. (C) Immunoblot analysis of chMyD88 in cell lysates of chicken DF1 cells transfected chSPOP treated with DMSO or 20mM MG132 for 6h. (D) and (E) Immunoblot of human MyD88 and mouse MyD88 in HEK293T cells and mouse embryo fibroblast cells.

SPOP is a highly conserved protein among different species, with only one amino acid difference between chickens and humans or mice (S4 Fig). To test whether the downregulation of SPOP on MyD88 is a common event among mammals, we constructed expression vectors of SPOP and MyD88 of human and mouse origin and transfected the plasmids into Hela cells and CHO cells. As expected, SPOP negatively regulated MyD88 in both human and mouse cells (Fig 2D and 2E), suggesting the highly conserved regulatory role of SPOP on MyD88.

### ChSPOP promotes K48-linked polyubiquitination of chMyD88

Protein ubiquitination is the first step in ubiquitin-proteasome-dependent protein degradation. SPOP is the substrate recognition adaptor of the SPOP–Cullin 3–RING box 1 ubiquitin ligase complex. Our above findings uncovered the importance of chSPOP in the regulation of chMyD88 degradation, so we next questioned whether chMyD88 was the authentic substrate of the chSPOP E3 ligase complex. To explore this possibility, chMyD88 and chSPOP were transfected into chicken cells in the presence of HA-tagged ubiquitin. Our results demonstrated that overexpression of chSPOP increased the ubiquitination level of chMyD88 in chickens (Fig 3A, lanes 1,2).

**Fig 3.**
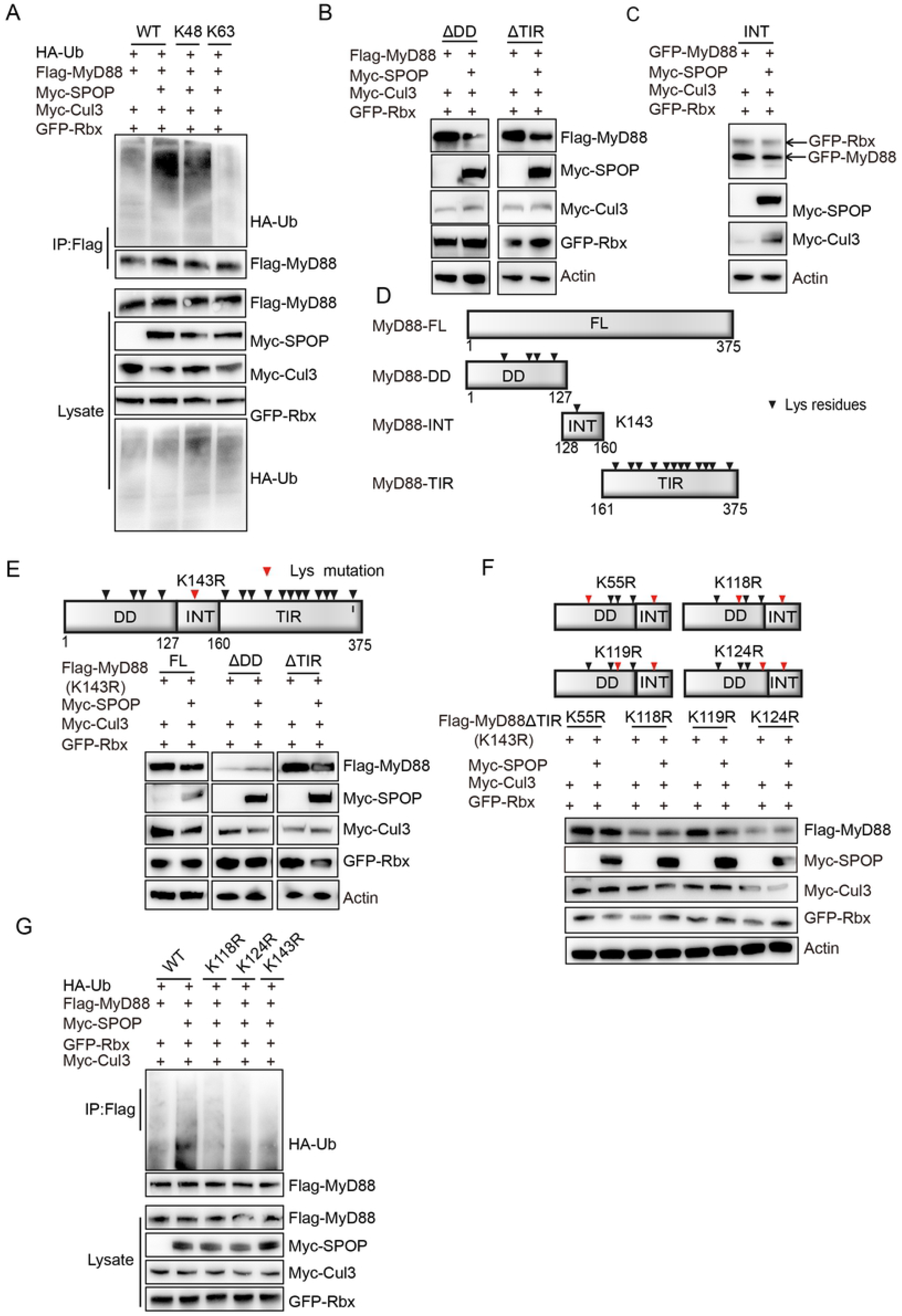
ChSPOP promotes K48 linked polyubiquitination of chMyD88. (A) Immunoblot analysis of immunoprecipitated chMyD88 from chicken DF1 cells transfected with Myc-tagged chSPOP with HA-tagged ubiquitin (HA-Ub), HA-tagged K48-linked ubiquitin (K48-Ub) or HA-tagged K63-linked ubiquitin (K63-Ub). Immunoprecipitation was carried out with anti-Flag antibody and probed with indicated antibodies. (B) Immunoblot analysis of lysates from chicken DF1 cells transfected with Flag-tagged chMyD88 truncations and Myc-tagged chSPOP. (C) Immunoblot analysis of lysates from chicken DF1 cells transfected with GFP-tagged intermediate domain of chMyD88 and Myc-tagged chSPOP. (D) Schematic diagram of the truncated chMyD88 mutants. (E) The Death-like domain of chMyD88 is required for the downregulation of chSPOP on chMyD88. Full length (K143R), K143R and Death-like domain truncated chMyD88, or K143R and TIR domain truncated chMyD88 was transfected with Myc-tagged chSPOP into chicken DF1 cells and cell lysates were immunoblotted with corresponding antibodies. (F) K118 and K124 of chMyD88 are required for the downregulation of chSPOP on chMyD88. K143R Death-like and intermediate domain of chMyD88 with K55R, K118R, K119R or K124R mutant was transfected into chicken DF1 cells. The expression of K143R Death-like and intermediate domain of chMyD88 was detected by anti-Flag antibody. (G) Mutations of K118, K124 and K143 reduce the polyubiquitination of chMyD88. Wild type, K118R, K124R or K143R chMyD88 was transfected into chicken DF1 cells with chSPOP. chMyD88 was immunoprecipitated by anti-Flag antibody followed by immunoblotting with indicated antibodies.

MyD88 could be ubiquitinated by either K48 or K63-linked ubiquitination[10–13]. To investigate the molecular mechanisms of SPOP-mediated degradation of MyD88, we transfected vectors expressing HA-tagged K48 or K63-linked ubiquitin into chicken DF1 cells, and found that overexpression of chSPOP enhanced the K48-linked rather than the K63-linked ubiquitination of chMyD88 (Fig 3A, lanes 3,4).

We next constructed a truncated form of chMyD88 to determine the ubiquitination domain of chMyD88. As the intermediate domain mediated the interaction between MyD88 and chSPOP, the death-like domain and the TIR domain deleted truncations, which both contained the intermediate domain were firstly transfected into chicken cells. Cell lysates were denatured and subjected to immunoblotting to examine the expression of truncated chMyD88. Our findings showed that chSPOP could lead to the downregulation of either the death-like domain or the TIR domain deleted truncation of chMyD88 (Fig 3B), indicating that the commonly conserved intermediate domain in both the death-like domain and the TIR domain deleted truncation of chMyD88 might be the ubiquitination target region of chMyD88. We thus transfected chSPOP with a sole intermediate truncation of chMyD88 into chicken cells, and as expected we observed a significant decrease in truncated chMyD88 (Fig 3C). indicating that the intermediate domain only would be able to initiate the degradation. There is only one lysine site in the intermediate domain (Fig 3D), we speculated that K143 might be a ubiquitination site of chMyD88. To investigate if there are other potential lysine sites in the Death-like domain or TIR domain, we mutated K143 in the intermediate domain into R143 and co-transfected SPOP with mutated full length or domain deleted MyD88 into chicken cells. We found K143R mutated full length chMyD88 could still be degraded by chSPOP (Fig 3E, lanes 1,2), suggesting there might be other ubiquitination sites besides K143. chSPOP could lead to the downregulation of TIR domain truncated K143R chMyD88 while chSPOP abolished such activity when co-transfected with Death-like truncated K143 chMyD88 (Fig 3E, lanes 3,4,5,6), these results clearly demonstrated ubiquitination sites in the Death-like rather than in the TIR domain. We then mutated the four lysine residues one by one in combination with K143R to check which mutant would rescue the downregulation of chSPOP on chMyD88, immunoblot analysis showed that K118R, K124R together with K143R, but not K119R, abolished the degradation (Fig 3F). Furthermore, we mutated K118, K124 and K143 into arginines and found that chSPOP failed to downregulate the mutated chMyD88 in the protein level (S5 Fig). Lastly, we transfected the K118R, K124R or K143R chMyD88 with chSPOP into chicken DF1 cells, immunoprecipitation results showed that all of the three mutations reduced the ubiquitination level of chMyD88 (Fig 3G). Taken together, these data suggest that chSPOP promotes the degradation of chMyD88 through ubiquitination on K118, K124 and K143 of chMyD88.

### ChSPOP enhances NF-κB activation and proinflammatory cytokine production in chicken macrophages

To assess the effect of chSPOP on proinflammatory responses in chicken macrophages, we treated chicken macrophage cells (HD11) with the TLR4 agonist LPS and measured the mRNA and protein levels of IL-1β and IL-8 as indicators of proinflammatory responses after chSPOP overexpression or knockdown. Our results showed that chSPOP overexpression efficiently decreased the production of IL-1β and IL-8 when macrophages were treated with LPS (Fig 4A and 4B and S6A Fig). To test the effect of endogenous chSPOP on the host immune response, we silenced the expression of chSPOP using short interfering RNA (siRNA). The expression of chSPOP was significantly decreased by 60% in macrophages transfected with chSPOP-specific siRNA. Knockdown of chSPOP induced the expression of IL-1β and IL-8 compared with controls (Fig 4C and S6B Fig). Moreover, an ELISA assay demonstrated that the production of IL-1β was significantly enhanced upon LPS stimulation when the expression of chSPOP was impaired (Fig 4D), suggesting that chSPOP inhibits proinflammatory responses in LPS-treated cells.

**Fig 4.**
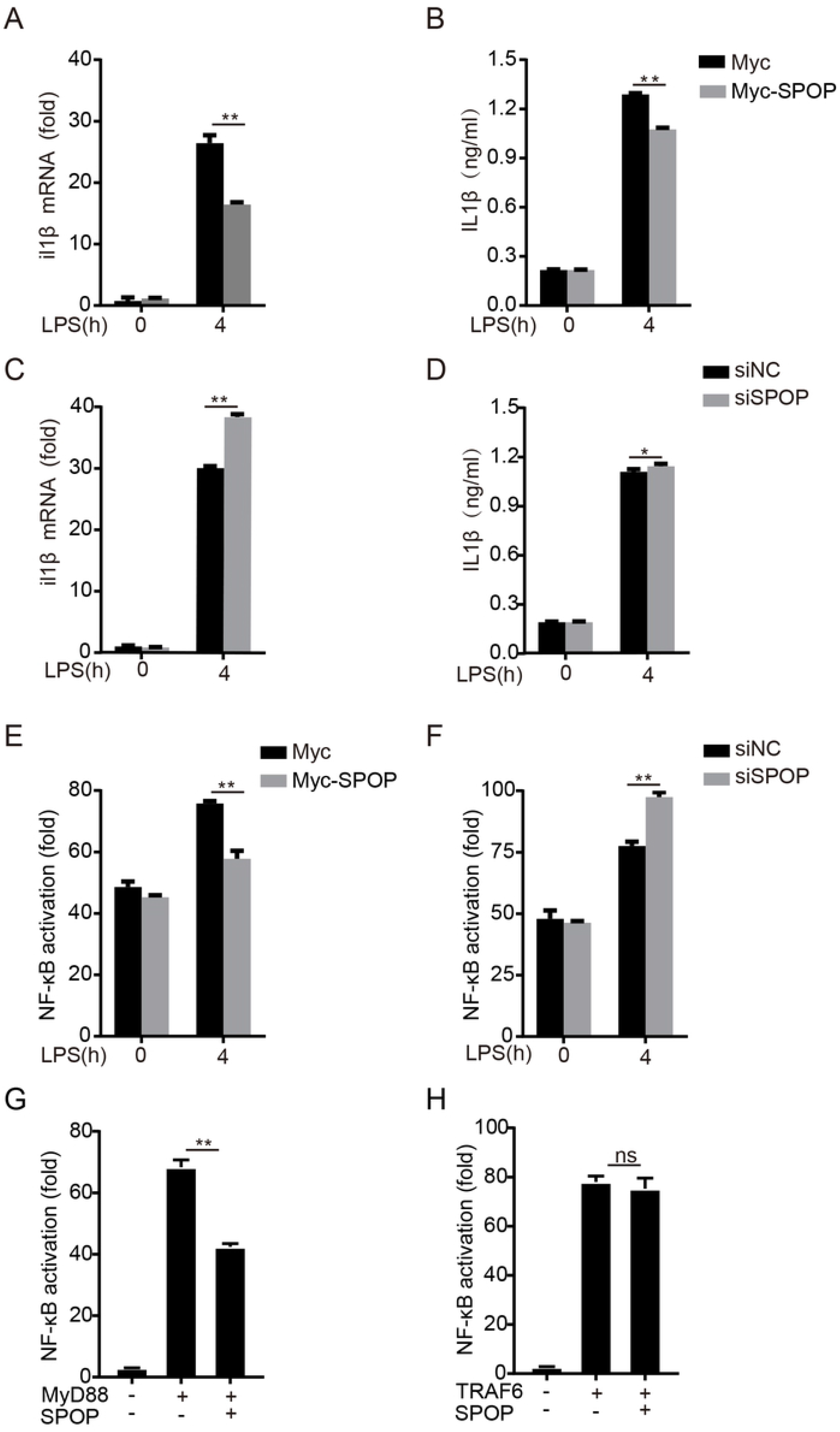
ChSPOP negatively regulates NF-κB signaling and IL-1β production. (A) and (B) The expression of IL-1β mRNA in mRNA and protein level of IL-1β in cell supernants of chicken HD11 macrophages of with overexpressed chSPOP and stimulated with LPS for 4 h. (C) and (D) The expression of IL-1β mRNA in cells and protein level of IL-1β in cell supernatants of chicken HD11 macrophages transfected with chSPOP siRNA and stimulated with LPS for 4 h. (E) Relative luciferase activity of NF-κB reporter in chSPOP overexpressed chicken HD11 macrophages. (F) Relative luciferase activity of NF-κB reporter in chicken HD11 macrophages transfected with chSPOP siRNA. These data are representative of three independent experiments. (G) and (H) Luciferase activity driven by NF-κB promoter in chicken DF1 cells transfected with SPOP and MyD88 or TRAF6. Luciferase assays were performed 24 h after transfection. Error bars reflect ±s.d.

To identify the molecular mechanisms through which chSPOP inhibits the LPS-triggered response, luciferase assays were performed to determine the effect of chSPOP on the NF-κB signal downstream of chMyD88. We expressed exogenous chSPOP or knockdown chSPOP by RNAi in chicken macrophage cells transfected with NF-κB reporter and then measured the NF-κB activity after LPS challenge. As expected, chSPOP negatively regulated LPS-induced NF-κB reporter activation (Fig 4E and 4F). We next assessed whether the manipulation of NF-κB signaling pathway of chSPOP was dependent on chMyD88 but not on other target proteins. In luciferase reporter assay, chSPOP overexpression inhibited chMyD88 mediated NF-κB activation (Fig 4G), while NF-κB activation mediated by overexpression of TRAF6 which was a downstream molecule of chMyD88 was not inhibited (Fig 4H). Taken together, our findings indicate the negative regulatory role of chSPOP on the MyD88-NF-κB signaling pathway and proinflammatory cytokine secretion.

### SPOP deficiency attenuates host defenses against *Salmonella* infection

To elucidate the *in vivo* function of SPOP, we generated SPOP^-/-^ conditional knockout mice using the Cre-LoxP recombination approach since germline knockdown of SPOP leaded to embryonic lethality. Wild-type and SPOP^-/-^ mice were injected intraperitoneally with *Salmonella typhimurium*, and the survival rates were monitored. SPOP^-/-^ mice were more susceptible to infection of *Salmonella typhimurium* (Fig 5A), and the knockout mice had about 10 folds more bacteria load in the spleen than did wild type mice after measuring the total colony-forming units (Fig 5B).

**Fig 5.**
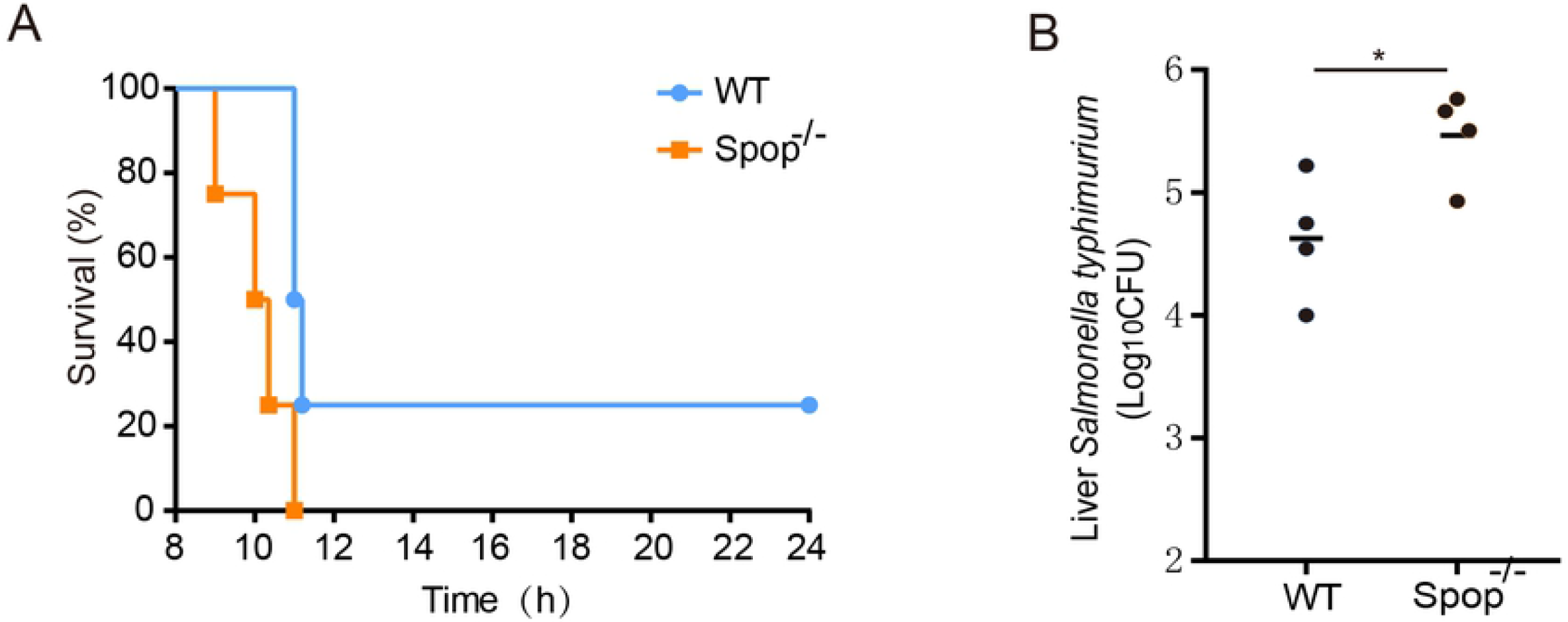
SPOP deficiency attenuates resistance against *Salmonella* infection in mice. SPOP^-/-^ heterozygous knockout mice were generated by Cre-LoxP recombination approach. Conditional knockout mice were feed with tamoxifen three times every 48h at the dose of 175μg/g body weight. (A) Mice were then challenged with *Salmonella typhimurium* and monitored the survival rates for 24 hours (n=4 each group). (B) Bacterial load in the liver of SPOP^-/-^ and wild type mice injected intraperitoneally with *Salmonella typhimurium*.

## Discussion

The recognition of bacterial PAMPs through cellular pattern recognition receptors triggers antibacterial responses to limit bacterial replication. In particular, MyD88-dependent TLRs are key pattern recognition receptors that can detect pathogen-derived LPS, flagellin or single-stranded RNA in the plasma membrane during infection with a variety of bacteria, such as Salmonella enteritidis and Escherichia coli. MyD88 has been identified as an essential adaptor protein for almost all TLR-dependent signaling [25] MyD88 is the point of convergence for sensing bacteria and DNA viruses, which then converts upstream signals to activate interleukin-1 receptor associated kinases (IRAKs), and ultimately leads to the production of proinflammatory cytokines [26]. Activation of MyD88-NF-κB signaling leads to antimicrobial responses to resist infection by pathogens. Meanwhile, inflammatory responses are dynamically modulated to maintain immune homeostasis through regulation of the half-live of the inflammation mediators and anti-inflammatory signaling. Excessive inflammation and the production of proinflammatory cytokines and interferons resulting from the misregulation of MyD88 signaling are detrimental to health and might lead to pathological damage, such as cancer and autoimmune diseases [27–29]. Several mechanisms that regulate the activity of MyD88 have been revealed [9–13, 18]. For example, transforming growth factor-β induces the Smad6-dependent recruitment of E3 ubiquitin ligases Smurf1 and Smurf2 to MyD88, targeting them for proteasomal degradation and thereby displaying its anti-inflammatory function [11]. E3 ubiquitin ligases Cbl-b and Nrdp1 are also described to polyubiquitinate and degrade MyD88 and inhibit TLR signaling to regulate antibacterial or antiviral responses [10, 12]. However, considering that different E3 ligases could be recruited to the same protein substrate and degraded through various mechanisms, more efforts are needed to demonstrate whether MyD88 can be targeted by other E3 ubiquitin ligases. Here, we uncovered a previously uncharacterized role of chSPOP in the regulation of chMyD88 protein abundance and the TLR signaling pathway and determined the underlying molecular mechanisms, using chickens as a model organism.

SPOP is an E3 ubiquitin ligase adaptor that is widely expressed in various organs. Emerging evidence has suggested that SPOP controls the stability of proteins involved in a range of cellular processes, such as tumorigenesis, senescence, transcriptional regulation and apoptosis [15, 16, 18–20]. Notably, SPOP has been extensively studied as a tumor suppressor and is a frequently mutated hotspot, most notably, in prostate cancer [16, 30, 31]. Cancer-associated SPOP mutants show reduced binding, ubiquitination and degradation of oncoprotein substrates, such as androgen receptor and ETS transcription factor ERG[16, 32]. In the current study, we characterized the distinct role of chSPOP on MyD88 and linked chSPOP to innate immune signaling. First, we showed that chSPOP promoted chMyD88 degradation by mediating K48-linked ubiquitination of chicken MyD88 Lys-118, Lys-124 and Lys143 residues. As expected, chSPOP recruited MyD88 to the Cullin 3-SPOP-RBX1 E3 ligase complex through its substrate binding MATH domain. To our knowledge, SPOP is the fourth E3 ligase to be identified that polyubiquitinates and degrades MyD88, adding to the complexity of the regulation of the MyD88-NF-κB innate immune signaling pathway. Second, we provide further evidence for the inhibition by chSPOP of chMyD88-induced NF-κB signaling and proinflammatory factors production. The possible effect of SPOP on NF-κB signaling is supported by a recent report showing that downregulation of SPOP promoted the migratory and invasive abilities of osteosarcoma cells by regulating the PI3K/Akt/NF-κB signaling pathway. Our findings expand the role of SPOP and uncover its association with innate immune signaling by modulating the adaptor MyD88. Third, we revealed the evolutionarily conserved mechanism of regulation of MyD88 by SPOP among birds and mammals. The amino acid sequence of SPOP is almost completely conserved, with only one amino acid substitution among humans, mice and chickens, while the sequence of MyD88 differs substantially between birds and mammals. However, we observed that both human and mouse SPOP interacts with and degrades MyD88.

Chickens have proven to be a versatile experimental model organism in the study of immunology, development biology, virology and cancer [33]. The current study firstly investigated the downregulation of chMyD88 by chSPOP in chickens, and then expanded the investigation into mouse and human cells. We generated a gene knockout mouse model to illustrate the in vivo function of SPOP. Of note, knockout of SPOP efficiently attenuates resistance to *S. typhimurium* infection in mice, further indicating the versatility of chickens as a model system in immunology.

In conclusion, we demonstrated that SPOP-mediated K48-linked ubiquitination and degradation of MyD88 through the proteasome pathway is a novel mechanism that negatively regulates MyD88-dependent proinflammatory signaling.

## Methods

### Ethics statement

Animal care and use protocols were performed in accordance with the regulations in the Guide for the Care and Use of Laboratory Animals issued by the Ministry of Science and Technology of the People’s Republic of China. The animal experiments were approved by the Animal Ethics Committee of the Institute of Animal Sciences, Chinese Academy of Agricultural Sciences (Approval Number: XK1917).

### Gene knockout mice and *Salmonella* infection

SPOP^-/-^ mice were created using classic Cre-LoxP recombination approach by Beijing Vitalstar Biotechnology Co., Ltd. SPOP^-/-^ mice were generated by crossing SPOP^-/-^ mice with Cre transgene C57BL/6 mice. Mice 8 weeks of age with similar body weight were kept in a specific pathogen free environment in China Agricultural University. SPOP^-/-^ mice and control mice were then challenged with *Salmonella typhimurium* (5*10^8^ CFU). For bacteria load analysis, liver samples were collected immediately after the death of mice.

### Cell culture and transfection

Human cervical carcinoma cells (Hela cells, from ATCC), Chinese hamster ovary cells (CHO cells, from ATCC) and DF1 cells (chicken embryonic fibroblast cell, from cell bank of Chinese Academy of Sciences) were cultured in Dulbecco’s modified Eagle’s medium supplemented with 10% fetal bovine serum (FBS, Gibco) and 1% penicillin-streptomycin (Gibco). chicken macrophage cells (HD11 cells) were cultured in RPMI1640 medium (Gibco) complemented with 10% FBS, 5% chicken serum, 1% sodium pyruvate, 1% non-essential amino acids and 1‰ β-mercaptoethanol. These cells were maintained in a humidified incubator with 5% CO_2_ at 37 °C. Lipofectamine 3000 (Invitrogen) was used for the transfection of plasmids or siRNA into HeLa, CHO and DF1 cells, according to the manufacturer’s instructions. Mirus purchased from Mirus Bio was used for the transfection of siRNA into HD11 cells. For certain experiments, cells were treated with MG132 (5 μM) for 4 h after transfection.

### Antibodies and reagents

The antibodies against FLAG, MYC and GFP (dilution ratio 1:2000) were purchased from Abmart. The polyclonal antibody against SPOP was purchased from Santa Cruz Biotechnology (1:500). Rabbit anti-MyD88 was purchased from Cell Signaling Technology (1:500). Mouse anti-β-actin antibody was obtained from Protocol (1:3000). The secondary HRP-conjugated antibodies used in this study were goat anti-mouse and goat anti-rabbit antibodies obtained from Abcam. LPS (Escherichia coli serotype O55:B4) was from Sigma-Aldrich.

### Plasmids

SPOP, MyD88, Rbx1 and Cul3 cDNAs were amplified using standard PCR techniques and high-fidelity DNA polymerase from a spleen cDNA library and were subsequently inserted into expression vector pcDNA3.1. Deletion mutants encoding different regions of the MyD88 or SPOP protein were obtained from full-length Flag-MyD88 or Myc-SPOP plasmids by PCR and were subcloned into pcDNA3.1. The Myc-tagged SPOP mutant (SPOP F133L) was obtained by two PCR amplifications using pcDNA3.1-Myc-SPOP as the template. HA-Ub, HA-UbiK48 (all lysines mutated to arginine except for K48) and HA-UbiK63 (all lysines mutated to arginine except for K63) were purchased from and were constructed using pBI-CMV. The lysine to arginine point mutants of MyD88 were generated using the QuikChange mutagenesis kit (Tiangen). All constructs were confirmed by sequencing. The NF-κB-Luc luciferase reporter plasmid was purchased from Promega.

### RNA interference

The siRNAs duplexes were synthesized by Gene-Pharma. The sequences of the siRNAs were as follows:

SPOP siRNA for chicken, 5’-GCCAGAACACUAUGAACAUTT-3’;
Nonspecific siRNA (N.C.), 5’-UUCUCCGAACGUGUCACGUTT-3’.

### Real-time PCR

Total cellular RNA was extracted by TRIzol (Invitrogen) according to the manufacturer’s instructions. Then, cDNA was generated from 1 μg of RNA using the PrimeScript RT reagent kit with gDNA Eraser (Takara). The mRNA quantifications of target genes were performed by real-time PCR using the SYBR GREEN MIX (Takara). Data were normalized to the expression of the housekeeping gene β-actin. The sequences of the PCR primers used to amplify the target genes are listed below:

β-actin: sense 5’-GAGAAATTGTGCGTGACATCA-3’,
antisense 5’-CCTGAACCTCTCATTGCCA-3’;
spop: sense 5’-AGGCTTGGATGAGGAGAGT-3’,
antisense 5’-CGCTGGCTCTCCATTGCTT-3’;
myd88: sense 5’-TGGAGGAGGACTGCAAGAAGT-3’,
antisense 5’-GCCCATCAGCTCTGAAGTCTT-3’;
il1β: sense 5’-GCATCAAGGGCTACAAGCTCT-3’,
antisense 5’-T CCAGGCGGTAGAAGATGAAG-3’;
il8: sense 5’-TCCTCCTGGTTTCAGCTGCT-3’,
antisense 5’-GTGGATGAACTTAGAATGAGTG-3’.

### Immunoprecipitation assay and immunoblot analysis

For the immunoprecipitation assay, cells transfected with the indicated plasmids were lysed in RIPA buffer (50 mM Tris-HCl, pH 7.4, 150 mM NaCl, 0.25% deoxycholic acid, 1% NP-40 and 0.5% SDS supplemented with protease inhibitor [Roche]) and centrifuged at 12,000 g at 4°C for 10 min. The whole cell lysate was precleared with protein A/G agarose and then incubated with anti-Flag beads or appropriate antibody and protein A/G agarose at 4°C overnight with constant rotation. Immunoprecipitated samples were collected by centrifugation and washed with RIPA buffer three times. After extensive washing, the immunoprecipitates were boiled in sample-loading buffer for 10 min to elute the precipitated proteins and subjected to immunoblot analysis.

For immunoblot analysis, the protein lysates or immunoprecipitate samples were separated on SDS-PAGE gels by electrophoresis and then transferred onto polyvinylidene fluoride membranes (Millipore). The membranes were first blocked with 5% (wt/vol) fat-free milk in TBST, then incubated with the corresponding primary antibodies diluted in 5% fat-free milk in TBST. After being washed with TBST, the membranes were incubated with the appropriate secondary antibodies diluted in fat-free milk in TBST. The protein bands were visualized by Immobilon Western Chemiluminescent HRP Substrate (Millipore) according to the manufacturer’s instructions.

### Luciferase reporter assay

Cells were seeded in 12-well culture plates and transfected with reporter gene plasmids (100 ng) combined with overexpression vector or siRNAs and other constructs as indicated. pTK-Renilla reporter plasmid was added to normalize the transfection efficiency. Twenty-four hours after transfection, LPS or sterile water was added to the cells (to a final concentration of 100 ng/ml). The luciferase activity was determined 4hr later using the Promega luciferase assay kit according to the manufacturer’s instructions.

### Measurement of cytokines

After transfection with the indicated plasmids or siRNA, cells were stimulated with 100 ng/mL LPS for 4 h. Then the culture supernatants were collected and the levels of the indicated cytokines were determined using a chicken IL-1B/IL-1 beta ELISA kit (LifeSpan BioSciences)

### Statistical analysis

Each experiment was repeated at least three times. Fold changes in mRNA levels (RT-qPCR), reporter assay activity and cytokine content between differently treated samples were compared using one-way ANOVA. In all analyses, P < 0.05 was considered statistically significant.

## Acknowledgments

We thank Beijing Vitalstar Biotechnology Co., Ltd for providing technical support in generation of conditional knockout mice of SPOP.

## Author contributions

Conceptualization, Q.L., F.W., J.W. and G.Z.; Methodology, Q.L., F.W. and Q.W.; Investigation, Q.L., F.W.; Writing–Original Draft, Q.L., G.Z. and F.W.; Writing – Review & Editing, Q.L., G.Z. and F.W.; Funding Acquisition, Q.W., R.L., M.Z. and H.C.; Resources, J.W. and G.Z.; Supervision, Q.L., J.W. and G.Z.

## Supporting information

**S1 Fig. Interaction of human and mouse MyD88 with SPOP.** HEK293T cells and mouse CHO cells were transfected with human (A) or mouse (B) MyD88 and SPOP. Immunoprecipitation were carried out to detect the interaction between MyD88 and SPOP by using the anti-FLAG antibody, followed by immunoblot analysis with indicated antibodies. Supporting information

**S2 Fig. Expression of chSPOP mRNA and chMyD88 in chSPOP overexpressed (A) and knockdown (B) DF1 cells.**

**S3 Fig. Immunoblot analysis of endogenous chMyD88 in chSPOP inhibited cells by CRISPRi.**

**S4 Fig. Comparison of SPOP amino acid sequences of human, mouse and chicken.**

**S5 Fig. ChSPOP failed to downregulated the protein level of K188, K124 and K143 mutated chMyD88.** Immunoblot analysis of chMyD88 in cell lysates of chicken DF1 cells transfected with chCUL3, chRBX1 and Myc-tagged chSPOP.

**S6 Fig. The expression of IL-8 mRNA in chicken HD11 macrophages of with overexpressed (A) or knockdown (B) chSPOP and stimulated with LPS for 4 h.**

